# Common to rare transfer learning (CORAL) enables inference and prediction for a quarter million rare Malagasy arthropods

**DOI:** 10.1101/2024.08.21.608960

**Authors:** Otso Ovaskainen, Steven Winter, Gleb Tikhonov, Nerea Abrego, Sten Anslan, Jeremy R. deWaard, Stephanie L. deWaard, Brian L. Fisher, Brendan Furneaux, Bess Hardwick, Deirdre Kerdraon, Mikko Pentinsaari, Dimby Raharinjanahary, Eric Tsiriniaina Rajoelison, Sujeevan Ratnasingham, Panu Somervuo, Jayme E. Sones, Evgeny V. Zakharov, Paul D. N. Hebert, Tomas Roslin, David Dunson

**Author notes:** shared first authorship (Otso Ovaskainen, Steven Winter).

## Abstract

Modern DNA-based biodiversity surveys result in massive-scale data, including up to millions of species – of which most are rare. Making the most of such data for inference and prediction requires modelling approaches that can relate species occurrences to environmental and spatial predictors, while incorporating information about their taxonomic or phylogenetic placement. Even if the scalability of joint species distribution models to large communities has greatly advanced, incorporating hundreds of thousands of species has not been feasible to date, leading to compromised analyses. Here we present a novel “common to rare transfer learning” approach (CORAL), based on borrowing information from the common species to enable statistically and computationally efficient modelling of both common and rare species. We illustrate that CORAL leads to much improved prediction and inference in the context of DNA metabarcoding data from Madagascar, comprising 255,188 arthropod species detected in 2874 samples.

## MAIN TEXT

Earth is home to several millions of species^1^. Among these, the majority are unknown^2^ and rare^3^. Recent innovations in sensor technologies are now providing unprecedented capacity to record patterns and changes in the abundance and distribution of all kinds of taxa, from the named to the previously unnamed and from the rare to the common. These technologies include DNA-based monitoring, passive acoustic monitoring, and visual sensors^4,5^. By allowing the efficient recording of thousands to hundreds of thousands of species in time and space, the accumulation of high-dimensional “novel community data” is transforming our access to information on species distributions and abundances^4^. As a particularly exciting development, the emergence of novel community data allows us to target the speciose groups accounting for the main part of global biodiversity^1,2^. Where species records to date have been massively biased towards vertebrates, one of the least species-rich taxa^3^, recent methods are now making hyper-diverse taxa such as arthropods and fungi arguably *easier* to sample than vertebrates and plants. As these speciose taxa can be mass-sampled and mass-identified, we can derive automated characterizations of what taxa occur where^5–7^. Nonetheless, the recent revolution in the generation of data is awaiting a matching insurgence of novel methods to analyze the data.

While most species on Earth are rare, these are the species that we know least about, partially because rare species are the most challenging to model^8^. Paradoxically, the rare species also encompass the taxa in greatest need of protection, and thus the very species for which information on their distributions and ecological requirements is most critical (‘the rare species paradox’^9^). Understanding biodiversity change necessitates models which can relate species occurrences to environmental, biotic, and spatial predictors, and which can predict changes in species communities with changes in the state of these drivers^10^. Hence, the need for predictive tools for rare species has been repeatedly highlighted^11–14^. However, the inherent rarity of most species results in highly skewed species-abundance distributions, where a few species are common whereas most species are present at few sites in low numbers. Typical approaches to species-level modelling will then impose a cut-off on species occurrences or abundances^15,16^ – arguing that for the rarest species, the data are simply insufficient for any quantitative inference regarding the drivers of their distribution. In a world where rare is common^3^, this can and will typically amount to rejecting most data, and all the information there then remains hidden. To make the most of increasingly available data, we need modelling approaches which can fully exploit such data.

With species distribution models (SDMs), rare species may be modelled through ensemble predictions from multiple small models, each of which contains just a few predictors to avoid overfitting^9,17,18^. Because closely related species are generally ecologically more similar than distantly related species^19,20^, phylogenetic information may be used to infer the distributions of rare species^21–25^. Joint species distribution models (JSDM)^26^ allow levelling up by modelling large numbers of species simultaneously. This enables efficient borrowing of information across species through their shared responses to environmental variation^27^. Furthermore, when data on species phylogenies and/or traits are available, information can be borrowed especially across similar species^28,29^. This can lead to improved predictions, especially for rare species^30^.

The high-dimensional, and often extremely sparse, nature of species occurrence data, compounded with spatiotemporal and phylogenetic dependencies, presents major challenges for statistical analyses and computation. High performance computing can scale some existing JSDMs to thousands of species^31,32^. Two-stage methods, which make small concessions by cutting dependence between species via approximate likelihoods^33,34^, can scale to tens of thousands of species while still retaining reasonable uncertainty estimates. Unfortunately, these approaches do not yet scale to the millions of species that comprise the Earth’s biodiversity^1^. What is more, they may perform poorly for extremely sparse rare species, by compromising model structures in the interest of gaining computational advantage.

## Results

### The Hierarchical Modelling of Species Communities framework

In this paper, we apply Bayesian transfer learning^35^ to develop the “common to rare transfer learning approach” (CORAL) (Fig. 1). Transfer learning refers to a broad class of multi-stage analysis methods which leverage information from a pre-trained model to improve performance for a new but related inference task. In a Bayesian context, this is often achieved by using the posterior model from one dataset to define an intelligent prior model for another dataset. Sharing information between models can improve parameter estimates and significantly boost out-of-sample performance, particularly when studying new, smaller datasets. Our transfer learning method builds on the Hierarchical Modelling of Species Communities (HMSC)^10,29,36^ approach to joint species distribution modelling (JSDM)^26^. The core idea of CORAL is very general and will thus apply to many other JSDM approaches, too. What makes its application in the HMSC context so intuitive is that HMSC models species responses to predictors as a function of species traits and phylogenetic relationships. This feature can be efficiently harnessed for transfer learning.

**Figure 1.**
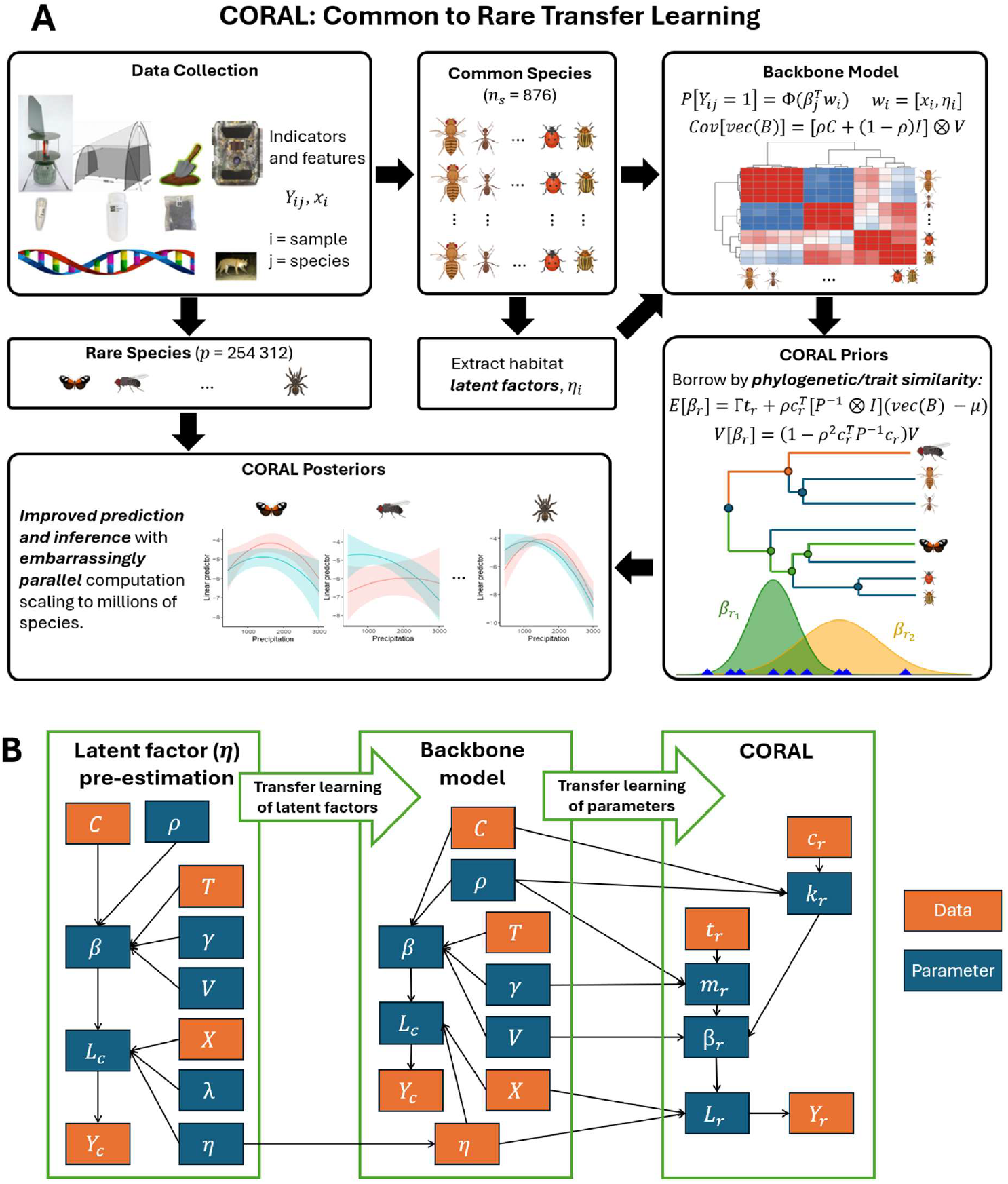
(A) High-level description of CORAL considering its application to Malagasy arthropods and (B) generic CORAL model structure implemented in the R package provided with this paper. (A) CORAL is based on fitting a backbone JSDM model to a subset of the most common species in the data and then borrowing information from this backbone to model the rare species. The backbone model learns about latent factors representing relevant missing environmental predictors, as well as about the species responses to both the measured and the latent predictors. The backbone model provides an informative prior distribution for each rare species. This is particularly efficient when we have access to phylogenetic or trait information, allowing information to be borrowed especially from common species closely related to the rare species, or species sharing similar traits with the rare species. As we show with a case study targeting a quarter million rare Malagasy arthropods, such informative prior distributions greatly improve the modelling of the rare species, both in terms of inference and prediction. (B) CORAL simplifies a fully Bayesian JDSM by replacing latent factors with a pre-estimated point estimate and by accounting for dependence from common to rare species – but not for dependence from rare to common species. For a full explanation of the data (orange squares) and the parameters (blue squares), see text and methods. The first panel shows the parameter and data dependencies used to fit HMSC to the common species only; this estimates the latent factors as a parameter (*η* in blue square), a point estimate of which is considered as data (*η* in orange square) in a second HMSC model fit to the common species. The second HMSC model, called the backbone model, has fewer parameters, significantly simplifying inference. Finally, parameters from the backbone model are used to define independent rare species models leveraging information from common species. To enable easy application of the CORAL approach to high-dimensional biodiversity data, we provide an R package for fitting these models, visualizing the results, and generating predictions.

In brief, HMSC is a multivariate generalized linear model fitted in a Bayesian framework. As a response it considers a matrix of species occurrences or abundances. We exemplify our approach with presence-absence data, denoting by *y*_ij_ = 1 if species *j* (with *j* = 1, …, *n*_s_) is present in sample *i* (with *i* = 1, …, *n*_y_) and *y*_ij_ = 0 if this is not the case. Presence-absence data are modelled in HMSC with probit regression: Pr(*y*_ij_ = 1) = Φ(*L*_ij_), where Φ(.) is the standard normal cumulative distribution function and *L*_ij_ is the linear predictor modelled as:

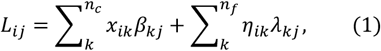

where *x*_ik_ are measured predictors, *η*_ik_ are latent predictors, and *β*_kj_ and *λ*_kj_ are regression coefficients quantifying responses of the species to the measured and the latent predictors. The latent features induce within-sample dependence across species; these features may encode characteristics of the habitat, the environment and the spatio-temporal setting not captured by the *x*_ik_s. HMSC uses a Bayesian hierarchical model to (a) automatically infer how many latent features *n*_f_ are needed and (b) to borrow information across species in inferring the *β*_kj_s. For (b) HMSC estimates to what degree taxonomically or phylogenetically related species, or species with similar traits, show similar responses *β*_kj_ to environmental variation through the multivariate normal distribution^10,28^

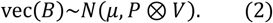

Here, *B* is the matrix of the regression parameters *β*_kj_ of the *p* species, *μ* is the average response, 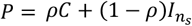 is a weighted average between the phylogenetic or taxonomic correlation matrix *C* and the identity matrix *I* corresponding to unrelated species, 0 ≤ *ρ* ≤ 1 is the strength of the phylogenetic signal, and *V* is the variance-covariance matrix of species-specific deviations from the average *μ*. The average response *μ* is further modelled as *μ* = vec(Γ*T*^T^), where *T* is a matrix of species traits, Γ are the estimated responses of the traits to environmental variation, and the superscript T denotes the matrix transpose. With Eq. 2, HMSC learns if and to what extent related species, or species with similar traits, show similar environmental responses. This allows for effective borrowing of information among species; for example, improving parameter estimation for rare species, for which it would be difficult to obtain accurate estimates if considering the data in isolation from the community context^29^. As a result, HMSC shows higher predictive performance compared to approaches that do not enable such borrowing of information^30^.

### Deriving conditional priors for rare species by borrowing information from common species

Our key idea is that even if it is not feasible to include 100,000+ species in a JSDM model such as HMSC, one can still borrow information from the common species. The structure of our approach follows naturally from three assumptions, namely that (1) users have enough data to perform high-quality inference on common species without leveraging rare species data, (2) information from these common species is relevant for modelling rare species, and (3) rare species may be viewed as conditionally independent given the common species data and measured sample covariates. This suggests a two-stage analysis which first studies the common species jointly and then studies each rare species independently using the results of the common species analysis.

The first stage of CORAL is to fit an HMSC to the common species *to pre-estimate latent factors* (Fig. 1). From this analysis we obtain a point estimate of the latent features 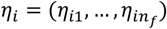, which provides key information not captured in 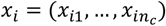 about environmental and habitat conditions and the overall biological community represented in sample *i*. We define a new covariate vector 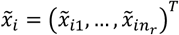 by concatenating *x*_i_ and *η*_i_ to be used as a fixed predictor in the second stage of CORAL, which fits HMSC to the common species to provide a *backbone model* (Fig. 1). The third stage fits CORAL models (independent Bayesian probit models) to each rare species: 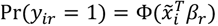, for *r* ∈ 𝒥_r_ with 𝒥_r_ ⊂ {1, …, *p*} the set of rare species (Fig. 1). To reduce mean square error in inferring *β*_r_ for *j* ∈ 𝒥_r_, we construct a prior which adaptively shrinks towards the common species coefficients accounting for taxonomic/phylogenetic similarity.

Our prior is motivated by the prior for fixed-effects coefficients in HMSC. To simplify inference and learn relevant hyper-parameters, we first re-run HMSC with the expanded covariate vector *x*-_i_. Under HMSC, the prior conditional distribution of the rare species coefficients given the common species coefficients is

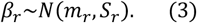

Here, the mean is given by

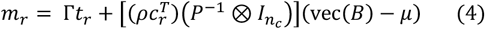

where *P* = *ρC* + (1 − *ρ*)*I*_p_. The variance-covariance matrix is *S* = *k*_r_*V*, where the variance scaling factor *k*_r_ is given by

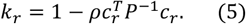

The vector *c*_r_ encodes relatedness between a rare species *r* ∈ 𝒥_r_ and all the *n*_s_ species in the backbone analysis and *t*_r_ is the trait vector for this rare species. As HMSC is fitted to data with Bayesian inference, parameter uncertainty can be accounted for by defining the prior as a mixture of multivariate normal distributions (each defined by Eqs. 3-5) over the posterior samples. To achieve a simple functional form for the prior, we approximated the mixture by a single multivariate normal distribution, the mean and variance-covariance matrix of which we set equal to those of the mixture (see Methods).

We refer to the above approach as <u>Co</u>mmon-to-<u>ra</u>re transfer learning (CORAL). Figure 1 shows the full mathematical structure of this approach, with each box corresponding to a separate stage of CORAL inference. This contrasts with the (computationally intractable) joint modelling approach, which would estimate all parameters for all species simultaneously. CORAL is likely to perform well when assumptions (1)-(3) hold: that is, when there is high-quality common species data that spans the phylogenetic tree and when the backbone model estimates that species responses are phylogenetically structured and/or influenced by species traits. To quantify the benefits of CORAL, we compare its performance to that of a baseline model that does not benefit from the backbone analysis. In other words, for the baseline model we fit 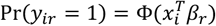, separately for *r* ∈ 𝒥_r_ using a simple Gaussian prior *β*_r_∼*N*(*m*_r_, *S*_r_). We expect CORAL to have substantial advantages over the baseline model due to two considerations: *x̃_i_* contains important latent factor information on top of *x*_i_, and CORAL allows the borrowing of information from the *β*_j_s for common species to rare species. As both CORAL and the baseline models can be fitted independently for *r* ∈ 𝒥_r_, computational time scales linearly with the number of species. As a result, these computations can be trivially parallelized allowing for inference and prediction for hundreds of thousands or even millions of species.

### Case study on Malagasy arthropods

We tested the approach in the context of metabarcoding data on Malagasy arthropods. We applied Malaise trap sampling in 53 locations across Madagascar, each of which was relatively undisturbed and where the vegetation represented the conditions of the local environment. We then applied high-throughput COI metabarcoding^37^ and the OptimOTU pipeline^38^ to score the occurrences of 255,188 species-level OTUs (henceforth, species) in 2874 samples (see *Methods*). To create a backbone model of common species, we included those 876 species that occurred in at least 50 samples. This left those 254,312 species that occurred less than 50 times in the data as rare species, which we model by the CORAL approach. We note that the threshold of 50 occurrences is relatively high so some of the rare species are not so rare. This choice was made to test the hypothesis that borrowing information from the backbone model changes predictions and inference especially for the very rare species – but less so for more common species. Most of these rare species were extremely rare in the sense that 182,402 species (71% of all rare species) were detected in one sample only. Among these extremely rare species, 1479 were singletons, i.e., represented by a single sequence. Some of these taxa may be artefacts, reflecting chimeric sequences or sequencing error. However, the vast majority (99.4%) of the rare species were represented by more than one sequence. Thus, the potential interpretation of some sequencing errors as false species is unlikely to qualitatively influence our conclusions.

As simple and frequently used predictors of species presence, we included covariates related to seasonality, climatic conditions, and sequencing effort. Climatic conditions were modelled through the second order polynomials of mean annual temperature and mean annual precipitation^39^, while including the interaction between these two climatic predictors. We modelled seasonality through periodic functions sin(2*πd*/365), cos(2*πd*/365), sin(4*πd*/365), and cos(4*πd*/365), where *d* is the day of sampling. To capture site- and sample-level variation not captured by the measured predictors, we included ten site-level (*n* = 53 sites) and four sample-level (*n* = 2874 samples) latent variables. Variation in sequencing effort was modelled by including log-transformed sequencing depth as a predictor. As a proxy of phylogeny, we used taxonomic assignments at the levels of kingdom, phylum, class, order, family, subfamily, tribe, genus and species, including assignments to pseudotaxa for those cases that could not be reliably classified to previously known taxa (see *Methods*).

The common species responded especially to site-level variation (Fig. 2A). This was shown both by responses to climatic variables, which contributed 48% of the explained variation, and by responses to the site-level random factors, which contributed 42% of the explained variation. The effects of the remaining predictors were much less pronounced, with seasonality contributing 3% of the explained variation, sample-level latent factors 7%, and sequencing depth 0.1%. As we did not include any traits in the model, we only based the CORAL models on borrowing information on taxonomic relatedness. The responses of the species to the predictors were strongly phylogenetically structured (posterior mean *ρ* = 0.65, posterior probability Pr(*ρ* > 0) = 1.00), thus providing potential for borrowing information especially from related species.

**Figure 2.**
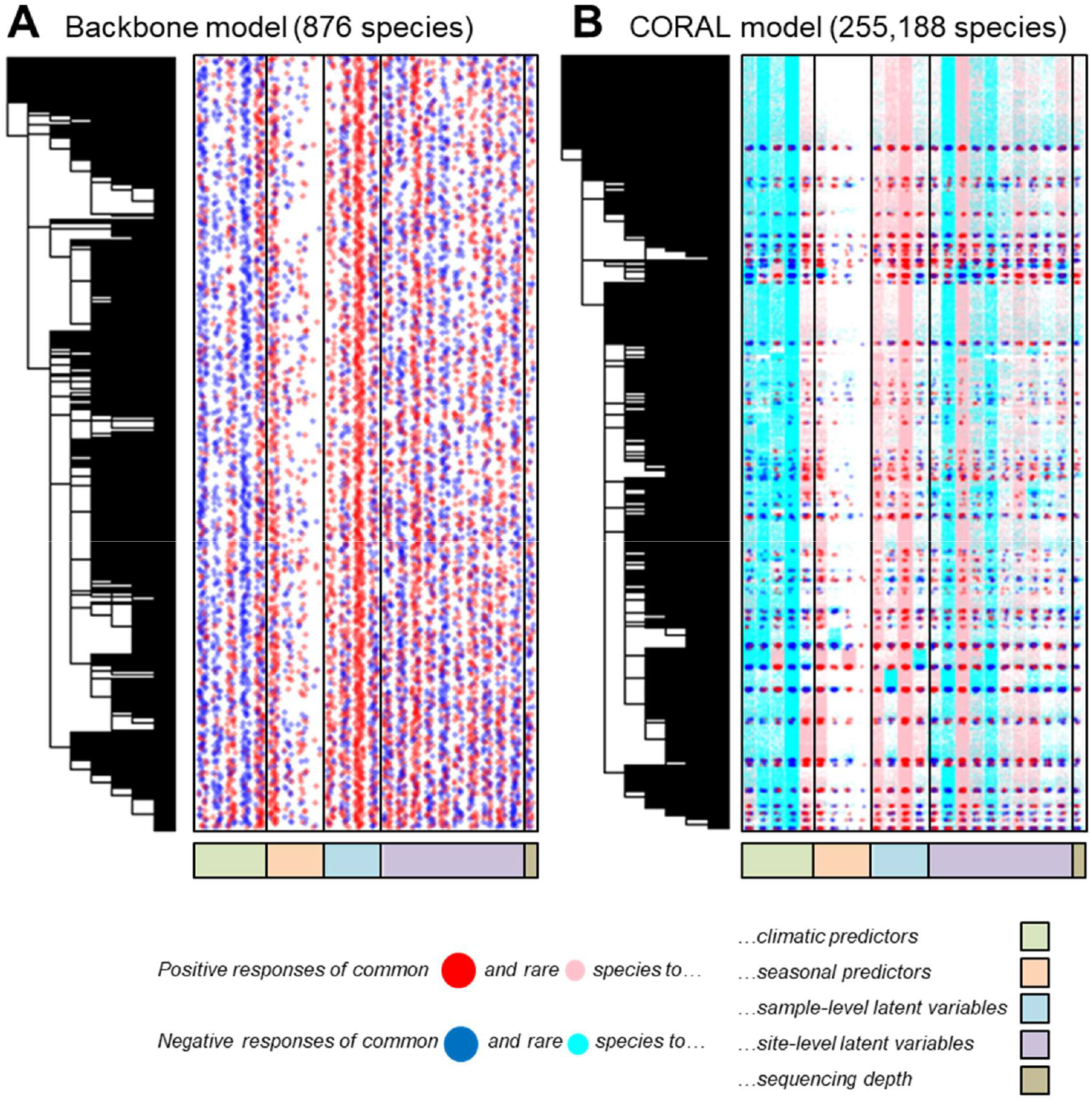
Responses of (A) the common species and (B) all species to measured and latent predictors. Responses that were estimated to be positive (red large dots for common species and pink small dots for rare species) or negative (blue large dots for the common species and cyan small dots for the rare species) with at least 95% posterior probability in the backbone model are highlighted. The dots have been made partially transparent and jittered in the horizontal direction to show the responses to many predictors for a very large number of species.

The variance scaling factor *k* varied between 0.13 and 0.70, with a mean value of 0.34, thus showing a substantial reduction in variance. As expected, it was lowest for species with close relatives in the backbone model (Fig. 3A). The conditional prior models predicted variation in the occurrences of the rare species better than random (Fig. 3B). This result is non-trivial as the predictions are made by a completely independent model that has not seen any data for the focal species. The accuracy of the prior predictions increased with the level of relatedness between the focal species and the species in the backbone model and the predictions were more accurate for species occurring at least 10 times in the data than for the very rare species (Fig. 3B).

**Figure 3.**
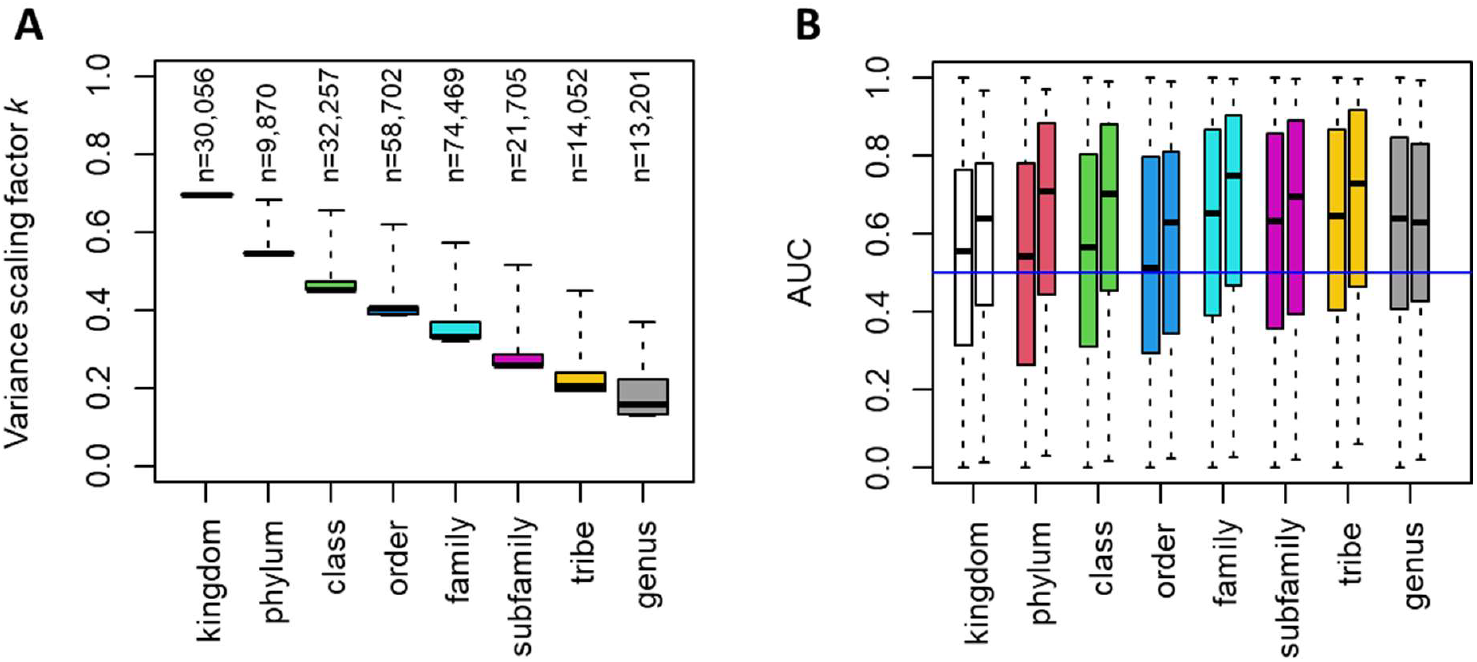
Conditional prior models for rare species constructed by borrowing information from the backbone model of common species. (A) Prior model precision measured by the variance scaling factor *k* (Eq. 5), as shown in relation to the taxonomic level shared with the closest relative in the backbone model. The numbers on top of the bars indicate the number of rare species in each category. (B) Discrimination powers of the conditional prior models, shown separately for each rank of the closest relative in the backbone model (different colors of bars) and for two prevalence classes (at most ten occurrences, left bars; more than ten occurrences, right bars). The blue line shows the null expectation AUC=0.5. In both panels, the lines show the medians, the boxes the lower and upper quartiles, and the whiskers the minimum and maximum values.

To compare the baseline and CORAL models in terms of inference, we fitted them to all of the 254,312 rare species. Combining the parameter estimates from the backbone and the CORAL models then enabled us to reveal the responses of all species (common and rare) to the measured and latent predictors (Fig. 2B). These responses illustrate how CORAL transfers information from common species to rare species, as in Fig. 2B blocks of red dots tend to spread pink dots in their surroundings, and blue dots tend to spread cyan dots in their surroundings, meaning that common species induce similar responses to taxonomically related rare species. However, there are exceptions to this general rule, as Fig. 2B shows the CORAL posteriors rather than the CORAL priors. Thus, if the data for a rare species has sufficient evidence of e.g. positive response even if the related common species show negative responses, the estimate of the rare species will be positive. By updating the conditional prior from the backbone model of common species with data from the focal rare species, we achieved improved predictions in the sense that the CORAL models showed higher precision than the baseline models (Fig. 4). This was especially the case for the very rare species (such as the one exemplified in Fig. 4B), for which the baseline models led to very large credible intervals, as may be expected for the low information contained in few occurrences. For more common species (such as the one exemplified in Fig. 4A), the increase in precision was smaller (Fig. 4C). The increase in precision increased with relatedness between the focal species and the species included in the backbone model (Fig. 4C), thus mirroring the relation seen between relatedness and the variance scaling factor (Fig. 3A).

**Figure 4.**
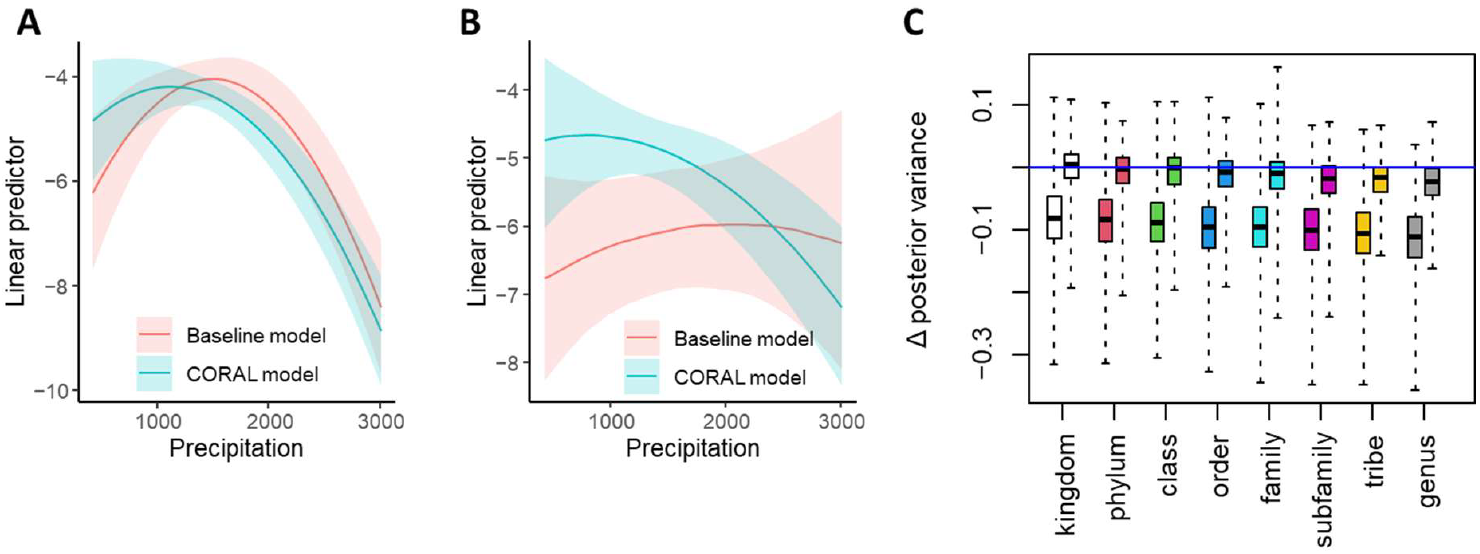
Comparison of inference between CORAL and baseline models. Panels A and B illustrate a specific prediction for two example species, one which is relatively common (A; the wall spider *Garcorops madagascar*, 10 occurrences) and another which is very rare (B; the deer fly *Chrysops madagascarensis*, 1 occurrence). The panels show the posterior mean (line) and the interquartile posterior range (shaded area) of the linear predictor under changing precipitation, keeping temperature at its mean value over the data. Panel C compares the posterior variance between baseline and CORAL models systematically for all species. We averaged the posterior variance over the environmental predictors (excluding intercept and latent factors). The panel shows the difference between posterior variance in the CORAL and baseline models. Thus, for values below 0 (the blue line) the CORAL model shows smaller variance. In panels C, the left-hand boxes correspond to very rare species (1-10 occurrences), the right-hand boxes to relatively common species (11-49 occurrences), the lines show the medians, the boxes the lower and upper quartiles, and the whiskers the minimum and maximum values.

One benefit of the CORAL approach is that its posterior distribution is presented analytically rather than through posterior samples obtained through MCMC. This is achieved by approximating the CORAL posterior for each species (both the common and the rare) by a multivariate normal distribution (see *Methods*). This saves storage space, which could otherwise become limiting for models with very large numbers of species. The multivariate normal presentation of the CORAL posterior also simplifies downstream analyses as posterior mean occurrence probabilities can be computed analytically without MCMC sampling (see *Methods*). The use of an analytical approximation may, however, introduce model misspecification, the extent of which we explored by comparing the posterior predictive distribution to the data in terms of relevant summary statistics (Fig. 5). The CORAL model fitted to the Malagasy arthropod data was well calibrated in terms of generally predicting the number of times each species was observed, except for some overestimation for the rarest species (Fig. 5A). The model also satisfactorily predicted the number of species present in each sample, but the overestimation in the occurrences of the rarest species translated to some overestimation of species richness (Fig. 5B). The model fit was uniform across ranges of temperature (Fig. 5C) and humidity (Fig. 5D), suggesting no substantial misspecification in terms of how the effects of these covariates were modelled.

**Figure 5.**
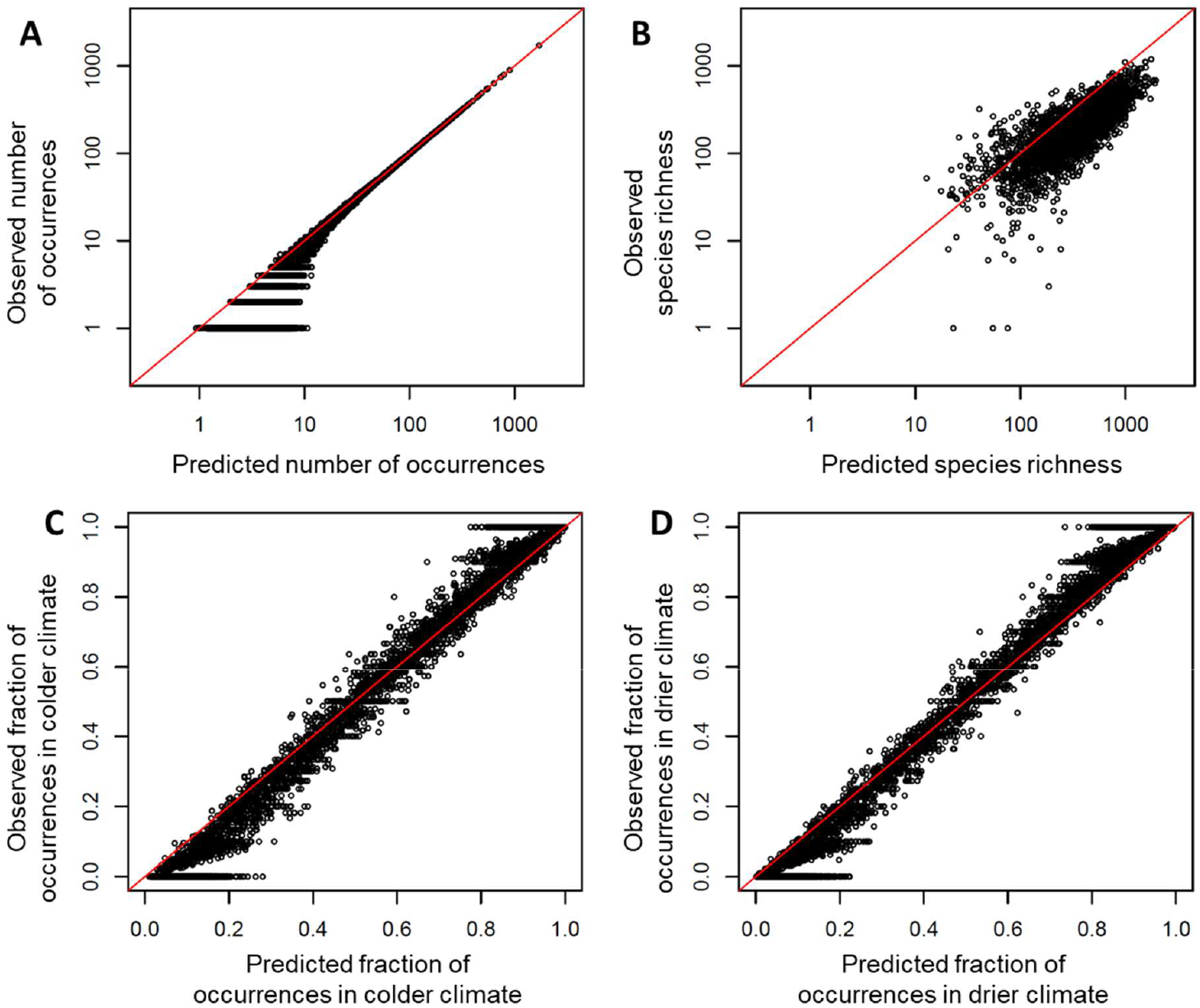
For inference to be trustworthy, it is key to verify that CORAL posteriors are consistent with the observed data, both in terms of the overall scale of the predicted probabilities as well as the learned covariate effects. (A) The expected number of observations for each species under the CORAL posterior compared to the observed prevalence, with the identity function is shown as a red line in all figures. CORAL probabilities are well calibrated in the sense that these predictions agree closely with the observed values, except for mild overestimation of the very rarest species. (B) Expected species richness for each sample under the CORAL posterior compared to the observed richness. Some overestimation is detectable among the very rarest species, which are roughly uniformly distributed across the samples. (C) and (D) Observed versus predicted proportion of occurrences below the median of (C) temperature and (D) precipitation, shown for species that occur at least 10 times in the data. CORAL probabilities are well calibrated across both covariates across their full range of values.

To compare the baseline and CORAL models in terms of predictive power, we considered the 22,140 species that were not included in the backbone model but occurred at least five times in the data. We applied two-fold cross validation, where we randomized the folds separately for each species, re-sampling until both folds included at least 40% of the occurrences. We compared the models using AUC, Tjur’s R^2^, PRAUC, Brier score, negative log-likelihood and log determinant posterior covariance. Together, these metrics provide a comprehensive overview of model performance covering predictive power, well calibrated probabilities, and useful inference (see *Methods* for their interpretations). All metrics of predictive performance improved considerably when moving from the baseline model to the CORAL model: AUC from 0.86 to 0.94, Tjur’s R^2^ from 0.03 to 0.08, PRAUC from 0.07 to 0.16, Brier score from 0.004 to 0.003, negative log-likelihood from 0.023 to 0.016, and log determinant from -28.2 to -36.4. All these improvements were significant with p<10^−16^ as measured by one-sided t-tests (see *Methods*). The improvement in the predictions was essentially independent of relatedness between the focal species and the species in the backbone model (Fig. 6), suggesting that most of the improvement derived from the inclusion of the latent factors estimated through the joint response of all common species, with less contribution from the direct borrowing of information from the related species. We validated this inference by fitting another set of models which included common species latent factors but not our novel prior; this approach retained about 75% of the gains in AUC over the baseline model. Additionally, the mean improvement in AUC did not essentially depend on the prevalence of the species (Fig. 6A), whereas for Tjur’s R^2^ the improvement was higher for the more common species (Fig. 6B).

**Figure 6.**
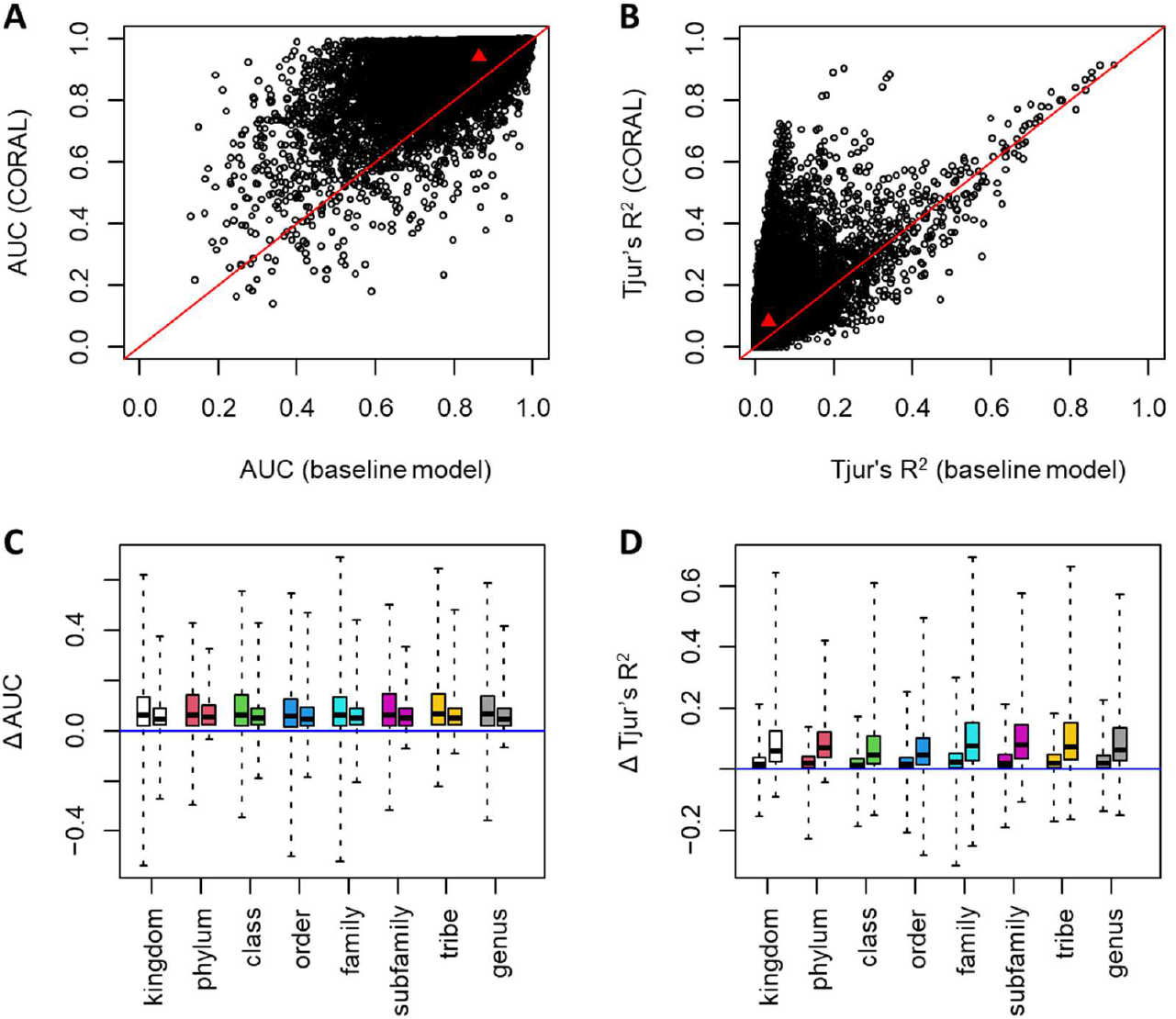
Comparison of predictive power for baseline and CORAL models based on two-fold cross validation. Predictive comparison is compared in terms of AUC (AC) and Tjur’s R^2^ (BD). The upper row of panels (AB) shows the raw values of the metrics for each species included in the analysis, with the red line showing the identity line and the red triangle showing mean values over the species. The lower row of panels (CD) shows the difference between the CORAL and baseline models. For values above 0 (the blue line), predictions by the CORAL model were more accurate. The results are shown separately for each rank of the closest relative in the backbone model (different colors of bars) and two prevalence classes (ten or fewer occurrences, left bars; more than ten occurrences, right bars). In panels C and D, the lines show the medians, the boxes the lower and upper quartiles, and the whiskers the minimum and maximum values.

## Discussion

The CORAL approach overcomes previous limitations on joint models of species communities with very large numbers of rare species. By borrowing information from a backbone model of common species, CORAL makes it possible to model even the rarest species in a statistically effective manner by combining an informative prior model with the limited data available for each rare species. As the rare species models can be parameterized independently, CORAL has an embarrassingly parallel implementation, making it feasible to analyze datasets comprised of millions of species. Rather than omitting rare species from all quantitative inference^15,16^, the approach developed here enables one to draw on the full information inherent in novel community data^4^. This allows one to generate informed predictions about changes in communities and overall biodiversity with changes in the state of environmental drivers. In essence, this amounts to putting the “-diversity” back in “biodiversity”.

We found species’ responses to climatic, seasonal, and latent predictors to be phylogenetically structured (posterior mean *ρ* = 0.65, posterior probability Pr(*ρ* > 0) = 1.00), forming the basis for borrowing information especially across related species. However, even without phylogenetic signal in the data, or alternatively by fitting a model without phylogeny, CORAL makes it possible to borrow information from the backbone model of common species by identifying sample- and site-level latent factors, as well as by basing the conditional mean on the average response of all species.

To illustrate the scale of the gain, we reiterate the proportion of rare species in our samples: had we imposed a cutoff of species occurrence in 50 samples, we would have omitted 254,312 out of 255,188 species (99.7%), retaining 876 species (0.3%) of the species pool. Leaving the rare species unmodelled would hardly be an efficient use of the massive data painstakingly acquired. For the 22,140 species (8.7%) which occurred at least five times in the data, but which were not included in the backbone model, we scored a substantial improvement in predictive power by borrowing information from the more common species. This is a major achievement, as it shows how the limited information inherent in the distribution of rare species may be leveraged by gleaning information from more common species.

While this study focused on methodological development, our findings are also of major interest for understanding the eco-evolutionary community assembly processes of the Malagasy fauna. We found seasonality and climatic responses of arthropods to vary with their phylogenetic relatedness, suggesting that their distributions across Madagascar are partially constrained by their ancestral niche. This region is characterized by extreme levels of endemism at both a regional and a very small scale^40–42^. Nonetheless, in adapting to local conditions, the species appear to maintain a strong signal of their ancestral niche.

## Acknowledgements

The work was funded by the European Research Council (ERC) under the European Union’s Horizon 2020 research and innovation programme (grant agreement No 856506: ERC-synergy project LIFEPLAN to OO, DD and TR). OO’s group was also funded by Academy of Finland (grant no. 336212 and 345110) and the European Union: HORIZON-CL6-2021-BIODIV-01 project 101059492 (Biodiversity Genomics Europe). JRdW, SLdW, MP, SR, JES, EVZ and PDNH were funded by grants awarded to PDNH from the Government of Canada through its New Frontiers in Research Fund (NFRFT-2020-00073), the Large Scale Applied Research Program administered by Genome Canada and Ontario Genomics (OGI-208), and the Major Science Initiatives Fund administered by Canada Foundation for Innovation (project number 42450). The authors wish to acknowledge CSC – IT Center for Science, Finland for computational resources.

## Author contributions

O. Ovaskainen^*^ acquired funding, coined the original idea, co-developed the statistical methodology, performed statistical analyses, contributed to software implementation, and co-wrote the first draft of the manuscript.

S. Winter^*^ coined the original idea, co-developed the statistical methodology, performed statistical analyses, contributed to software implementation, and co-wrote the first draft of the manuscript.

G. Tikhonov co-developed the statistical methodology, led software implementation and contributed to the first draft of the manuscript.

N. Abrego contributed to first draft of the manuscript, in particular placing the statistical framework into ecological context.

S. Anslan contributed to the implementation of the bioinformatics workflow.

J. R. deWaard contributed to sample management, DNA extraction and sequencing, and commented on the manuscript.

S. L. deWaard contributed to sample management, DNA extraction and sequencing.

B. L. Fisher acquired funding, participated in project coordination, participated in data collection and commented on the manuscript.

B. Furneaux led the implementation of the bioinformatics workflow and commented on the manuscript.

B. Hardwick participated in project coordination, participated in data collection and commented on the manuscript.

D. Kerdraon participated in project coordination and participated in data collection.

M. Pentinsaari contributed to the implementation of the probabilistic taxonomic classification method utilized in the bioinformatics pipeline.

D. Raharinjanahary participated in project coordination and participated in data collection.

E. T. Rajoelison participated in project coordination and participated in data collection.

S. Ratnasingham contributed to sample management, DNA extraction and sequencing.

P. Somervuo led the implementation of the probabilistic taxonomic classification method utilized in the bioinformatics pipeline and commented on the manuscript.

J. E. Sones contributed to sample management, DNA extraction and sequencing.

E. V. Zakharov contributed to sample management, DNA extraction and sequencing.

P. D. N. Hebert contributed to sample management, DNA extraction and sequencing, and commented on the manuscript.

T. Roslin contributed to first draft of the manuscript, in particular placing the statistical framework into ecological context.

D. Dunson acquired the funding, coined the original idea, co-developed the statistical methodology, and contributed to the first draft of the manuscript.

*Equal contributions

## Data availability

The data needed to reproduce the analyses are provided at Zenodo: https://doi.org/10.5281/zenodo.11076832. All raw sequence data are archived on mBRAVE and are publicly available in the NCBI Sequence Read Archive (SRA; www.ncbi.nlm.nih.gov/sra/SRA; BioProject accession TBD).

## Code availability

The R-based CORAL pipeline needed to reproduce the analyses is provided at Zenodo: https://doi.org/10.5281/zenodo.11076832. The bioinformatics pipeline is available at https://github.com/brendanf/CORAL_bioinfo.

## Inclusion & Ethics

The case study on Malagasy arthropods included local researchers (D. Raharinjanahary and E. T. Rajoelison) and was conducted in collaboration with a local partner (Madagascar Biodiversity Center). The roles and responsibilities amongst collaborators were agreed ahead of the research through an Access and Benefit Sharing Agreement.

## METHODS

### Deriving the CORAL Prior

CORAL is motivated by the default prior for coefficients in HMSC. Under this prior, the prior for a species *r* that is not part of the backbone model (i.e., a rare species) is given by *β*_r_ | *B* ∼ *N*(*m*_r_, *S*_r_), with the mean (4) and variance (5) given by the conditional multivariate normal formulas. The moments of this distribution are functions of HMSC parameters including Γ, *ρ, V*, and *B* and do not include information from the common species data, *Y*_c_, a-priori. Fitting the backbone model produces a posterior distribution, π, over these parameters, which in turn implies a posterior marginal distribution for the rare species coefficients,

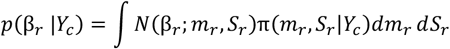

This updated distribution is our desired rare species prior. As this distribution is analytically intractable due to the integral over the posterior; we approximate it with a Gaussian:

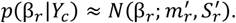

The mean 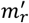 and variance 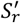 of this Gaussian are chosen to be the mean and variance of *p*(*β* |*Y*), respectively. These can be calculated using the laws of total expectation/variance, resulting in simple expressions in terms of posterior means/variances: 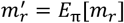 and 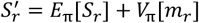. In practice, we approximate posterior means/variances using Monte Carlo with posterior samples returned by HMSC. This completes specification of the CORAL prior.

### Computational details

We fitted the backbone model with a high-performance computing accelerated version^32^ of the R-package Hmsc^36^, sampling each of the four chains for 37,500 iterations. Of these chains, we omitted the first 12,500 iterations as transient and then thinned the remaining chains by 100 to obtain 250 samples per chain and thus 1000 posterior samples in total.

For each rare species, we fitted a single-species model where we either did not (the baseline model) or did (the CORAL model) utilize information from the backbone models of common species. The baseline models were simple probit models with a Gaussian prior on the regression coefficients. The baseline models did not include the latent factors as predictors, and they assumed a default prior distribution for the species responses (N(0, 10) for the intercept and N(0, 1) for fixed effect coefficients). In the CORAL models, we included the latent factors as predictors, and assumed the conditional prior distribution based on Eqs. 3-5. We obtained 5000 samples after 2500 transient iterations for each species for both the baseline and CORAL models using MCMCpack^43^.

For each species, we summarized the CORAL model in terms of the mean *μ* and variance-covariance matrix Σ of the posterior samples. As the model contained 25 parameters (including the intercept), the model for each species was thus represented by 25 + 25(25 + 1)/2 = 350 parameters (accounting for the symmetry of Σ). The collection of models for all the 255,188 species thus contained ca. 89 million parameters, which resulted in the manageable file size of ca. 1.1 GB. We approximate the CORAL posterior through the multivariate normal distribution *N*(*μ*, Σ). For predictor vector *x*_i_, the posterior mean of the linear predictor can be then computed as 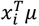, and the posterior mean of the occurrence probability as 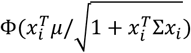.

### Metrics used to evaluate model performance

AUC is the probability a randomly chosen positive sample has a higher predicted probability than a randomly chosen negative sample. Tjur’s R^2^ is a pseudo-R^2^ value which can be read like any other R^2^ value, but typically reaches lower values^44^. PRAUC is the area under the precision recall curve – this metric quantifies true positives and is useful for analyzing highly imbalanced data where the minority class is of primary interest. The Brier score is the average squared error between predicted probabilities and labels – this metric penalizes overconfidence. Negative log-likelihood directly measures goodness of fit under the proposed Bernoulli model. The determinant of the posterior covariance determines the volume of a 95% credible interval for fixed effect coefficients under a Gaussian approximation, smaller intervals meaning more confident inference. Together, these metrics provide a detailed summary of discrimination ability (AUC, PRAUC), confidence (R^2^, Brier score), goodness of fit (negative log-likelihood), and precision (log-determinant).

### Sampling of Malagasy arthropods

The sampling was conducted as part of the worldwide LIFEPLAN biodiversity sampling design^45^. We selected 53 locations across Madagascar that were relatively undisturbed and where the vegetation represents the conditions of the local environment. 28 of the sites were sampled in a spatially nested sampling design with decreasing distances between them (50 km, 5 km and 500 m apart). The other 25 sites were spread across different forested habitats in Madagascar (dry, lowland and montane forests), at elevations ranging from 8 to 1592 MASL. We continuously collected one-week samples of flying arthropods in 95% ethanol using Malaise traps (ez-Malaise Trap, MegaView Science Co.) For a detailed description of the sampling, sample shipping and handling, and steps related to DNA extraction and sequencing, we refer to the LIFEPLAN Malaise sample metabarcoding protocol^46^.

The CORAL method can be applied to sample x species occurrence data generated by a wide variety of detection technologies and analysis pipelines. In this paper, we applied CORAL to DNA sequence data analyzed using the OptimOTU bioinformatics pipeline^38^, which was originally developed for the Global Spore Sampling Project^47^ and updated to apply to arthropod COI sequence data as part of the LIFEPLAN biodiversity sampling project^45^. The OptimOTU workflow for COI sequence data consists of primer removal, quality filtering, denoising, de novo and reference-based chimera removal, flagging likely non-animal sequences, removal of putative nuclear-mitochondrial pseudogenes, probabilistic taxonomic assignment, and finally taxonomically-guided hierarchical clustering. The OptimOTU pipeline is implemented using the targets 1.5.1 workflow management package^48^, here executed using the crew 0.9.0^49^ and crew.cluster 0.3.0^50^ backends in R 4.2.3^51^ on the Puhti cluster at CSC – IT Center For Science, Finland. This yielded a full taxonomic tree with approximate placeholder taxa to group those sequences which could not be reliably identified.

## Notes

https://doi.org/10.5281/zenodo.11076832

### Competing Interest Statement

The authors have declared no competing interest.

### Summary of Updates

We have revised the text for clarity, added new illustrations, and made the software more user friendly.

